# Beyond Deficit and Coexistence: Modeling the Knowledge–Conspiracy–Mistrust Configuration in Public Understanding of Science

**DOI:** 10.64898/2026.01.16.699843

**Authors:** Ahmet Süerdem, Svetlomir Zdravkov, Martin J. Ivanov

## Abstract

Debates about public trust in science often contrast deficit-based models, which emphasize the role of scientific knowledge, with constructivist perspectives that highlight the coexistence of multiple epistemologies. However, both approaches tend to overlook the mechanisms that link scientific knowledge, alternative epistemic orientations, and mistrust in science. To address this gap, the study applies a multilevel structural equation model within a multidimensional framework to examine conspiratorial reasoning as a key mechanism through which scientific knowledge influences science mistrust. Using cross-national survey data from Europe during the COVID-19 pandemic, the analysis also considers how this pathway is moderated by individual cognitive, motivational, and ideological traits, as well as macro-level political, cultural and economic factors. The findings reveal that conspiratorial reasoning significantly mediates the relationship between scientific knowledge and mistrust at both individual and regional levels. Moreover, the strength of these associations is conditioned by factors like informational engagement, regional value climates, and religiosity. Overall, the results suggest that scientific knowledge serves as a conditional epistemic resource, rather than a consistent buffer against mistrust in science.

## I. Introduction

Over decades of research, the concept of Civic Scientific Literacy (CSL) has been developed to capture the minimum level of scientific understanding required for citizens to engage meaningfully in democratic debates about science and technology [1–3]. Basically, CSL comprises four cognitive-attitudinal components - content knowledge, procedural understanding, appreciation of the positive impact of science, and the rejection of superstition. The framework conceptualized these components as tightly aligned and measurable along a single unidimensional scale. This approach was subsequently adopted by major international survey programs, establishing a standardized metric for assessing public understanding of science across nations.

Such unidimensional conception reflects the logic of the Deficit Model (DM), which attributes public mistrust of science to ignorance and assumes that increasing knowledge automatically produces acceptance of scientific institutions. However, this linear view has been widely criticized for ignoring the social and cultural contexts shaping public understanding of science. Wynne [4] showed that lay mistrust often arises not from ignorance but from experiences of institutional unresponsiveness and marginalization, while Jasanoff [5] critiqued technocratic models that privilege elite knowledge and dismiss lay reasoning. These critiques led to Public Engagement with Science (PES) approaches, which stress that perceived ignorance often reflects contextually grounded understanding shaped by identity, values, and lived experience [6].

While PES offered an important corrective to the DM’s reductionism and promoted more participatory science–society relations, it has also generated methodological and analytical limitations. As Irwin [7] observed, the field has fragmented into small-scale qualitative studies that, despite rich local insight, struggle to produce systematic or generalizable patterns. This parochial focus poses two main risks: overlooking structural and cross-cultural factors that shape public engagement, and blurring the boundary between expert and lay knowledge [8]. Such conflation complicates the evaluation of validity claims [9] and risks epistemic relativism, where all forms of knowledge are treated as equally authoritative.

Consequently, both the DM and PES share a critical blind spot: neither satisfactorily theorizes how individual components of CSL interact with contextual conditions to shape public understanding of science. PES rightly foregrounds local identities and lived experience, but often conflates individual agency with structural context, making it difficult to disentangle distinct psychological and social mechanisms. In contrast, the DM offers clearer constructs and causal pathways but does so at the cost of analytical reductionism. These limitations highlight the need for an integrative framework that treats CSL as multidimensional and examines how its components interact with both individual cognitive and motivational factors and broader macro-structural contexts.

Recent scholarship has increasingly adopted integrative perspectives on the relationship between scientific knowledge and attitudes, a direction to which this study contributes. While prior research has focused on direct associations between the two, systematic investigation of the indirect mechanisms remains limited. To address this gap, the present study investigates both individual- and macro-level pathways connecting key dimensions of CSL - namely, knowledge, attitudes, and superstition, operationalized here as conspiracy reasoning.

In what follows, we develop a theoretical framework that reconceptualizes CSL as a multidimensional, context-dependent configuration, emphasizing the mediating role of conspiratorial reasoning in linking scientific knowledge to institutional mistrust. We then outline a multilevel analytical model and derive testable hypotheses capturing both individual- and regional-level pathways. Drawing on cross-national survey data collected across Europe during the COVID-19 pandemic—a period marked by acute epistemic uncertainty and contestation of institutional authority—we examine these relationships using multilevel structural equation modeling. The paper concludes by considering the implications of the findings for broader debates on public understanding of science, epistemic authority, and science–society relations, moving beyond deficit-based and relativist accounts.

## II. Previous research and theoretical consideration

### A. Beyond Simple Deficits: Revisiting Civic Scientific Literacy Components

Recent scholarship increasingly transcends the rigid DM–PES divide. Rather than rejecting deficit-based explanations, scholars now seek to integrate core CSL constructs with context-sensitive models of public engagement. Einsiedel [10], for instance, emphasizes the existence of “multiple publics” and the need to situate CSL within broader social contexts. Methodological and empirical work further shows that quantitative approaches need not be tied to linear deficit assumptions and that attitudes reflect interactions among knowledge, socio-cultural orientations, and institutional contexts [11,12]. Collectively, these contributions have steered the field toward “theories of the middle range” [13], which capture the multidimensional and multilevel character of science–society relations while retaining key explanatory variables.

Within this synthesized framework, knowledge still remains an important predictor of attitudes, retaining independent effects even as contextual factors condition its influence [11,14]. Large-scale multilevel research supports a modest but consistent association between knowledge and favorable attitudes across diverse national contexts [15]. However, this general relationship is not uniform: in contentious contexts, the link often weakens or even reverses. Knowledge’s protective effect appears contingent on the issue at hand, proving less effective in areas characterized by moral controversy or heightened perceived risk [16–18]. For instance, while knowledge may predict positive attitudes toward vaccines and GMOs, it is less consistently related to climate change skepticism or institutional trust [19]. When scientific claims threaten core values or group identities, motivated reasoning can lead to increased polarization, with higher knowledge sometimes aligning with stronger resistance through identity-protective cognition [20].

Furthermore, CSL encompasses not only knowledge and attitudes but also the rejection of superstition, broadly understood to include conspiratorial, pseudoscientific, and denialist epistemologies. Within the literature, these epistemic orientations are frequently described as “unwarranted epistemologies” or forms of “contaminated mindware” [21,22], denoting cognitive and psychological deficits that distort evidence evaluation rather than coherent alternative ways of reasoning. They are typically framed as the antithesis of rational thought, associated with a dogmatic cognitive style characterized by lower openness to experience, resistance to disconfirming evidence, and limited reflective reasoning [23]. Moreover, endorsement of one such belief system often predicts endorsement of others - a pattern labeled monological reasoning, in which interrelated beliefs dogmatically reinforce one another, sustaining a unified but distorted worldview [24,25].

Among these belief systems, the link between conspiratorial thinking and rational reasoning has received substantial scholarly attention in recent years. Lewandowsky et al. [26] found that while motivational factors such as political ideology predict anti-scientific attitudes in domain-specific ways, conspiracy belief consistently emerges as a strong, cross-domain predictor. It has also been associated with deficits in information processing, including increased susceptibility to misinformation [27], as well as with psychological traits such as a heightened need for control and certainty, and, in some cases, paranoia or delusion-like ideation [28–29]. Accordingly, stronger endorsement of conspiratorial beliefs is frequently linked to lower educational attainment, reduced cognitive complexity, weaker analytic reasoning, and broader tendencies toward irrationality [23,29,30].

Although labeling a claim as a *“conspiracy theory”* imposes the stigma of *“crippled epistemology”* [31], effectively marking it as distorted or irrational, empirical evidence shows that conspiratorial beliefs are widespread and not confined to pathological or fringe groups [32]. Meta-analytic reviews reveal only weak or inconsistent associations with stable personality traits once methodological differences are controlled [33,34]. In light of these findings, scholars increasingly conceptualize conspiracy beliefs as counter-authoritative epistemologies [35], interpreting them as culturally and epistemically embedded responses to uncertainty rather than as unequivocal indicators of cognitive or psychological dysfunction [36–38].

Research increasingly highlights the contingent relationship between scientific and alternative epistemologies. In Japan [39] and Taiwan [40], for instance, higher levels of factual knowledge have been shown to positively predict engagement with pseudoscientific practices - patterns attributed to intellectual curiosity and dialectical cognitive orientations that favor reconciling apparent contradictions. In the Nigerian context, Falade and Bauer [41] interpreted the interaction between religiosity and knowledge in predicting attitudes towards science as suggesting that scientific and religious epistemologies can coexist, with scientific understanding not necessarily contradicting religiosity across its varying intensities.

### B. Epistemic Pluralism Without Relativism: On Coexistence, Conflict, and the Conditions of Justification

The coexistence of scientific and lay epistemologies is often explained through cognitive polyphasia (CP), which holds that individuals draw on multiple, culturally embedded epistemic frameworks to interpret and legitimize truth claims [42]. From this perspective, commonsense knowledge does not merely oppose science but provides shared meanings that shape identity and help manage uncertainty. Though seemingly irrational to outsiders, such reasoning remains psychologically coherent and socially meaningful to those who employ it [43]. These lay forms of knowing can also resist or reframe scientific meanings, influencing both research agendas and the translation of science into practice [44]. CP thus portrays knowledge production as a dialogical process in which scientific and non-scientific epistemologies coexist and mutually influence one another.

Despite its heuristic appeal, CP faces significant conceptual and methodological limitations if not carefully operationalized. Conceptually, its ambiguous formulation risks becoming a catch-all for epistemic diversity rather than serving as a precise analytical framework for understanding how competing truth claims coexist—or come into conflict. Methodologically, CP is challenged by its discursive and context-sensitive nature, often requiring in-depth qualitative fieldwork and resisting straightforward quantitative operationalization, especially across cultural contexts [45]. This necessitates mixed-methods approaches; however, without rigorous integration, such combinations risk degenerating into ad hoc eclecticism characterized by superficial ethnographic insights and unprincipled statistical interpretations, ultimately undermining both validity and accountability [46]. Overreliance on simplistic quantitative analysis to demonstrate the coexistence of science and religion (or superstition) can obscure the nuanced, contingent dynamics of epistemic interaction, conflating abstract constructs with the situated practices through which individuals negotiate meaning.

Moreover, CP must contend with the uneven power dynamics that shape how truth claims are legitimized and contested. Knowledge is embedded in power relations that influence the reconstruction, recognition and distribution of epistemic authority [46,47]. Any application of CP must therefore go beyond acknowledging plural perspectives to critically interrogate the conditions under which epistemic legitimacy is constructed and maintained. Blurring the boundary between lay and scientific epistemologies can be problematic, as both are internally fragmented and composed of multiple, sometimes conflicting, configurations. Apparent harmonies may reflect only transient alignments produced by contextual asymmetries.

In such fluid epistemic landscapes, justification becomes precarious, with standards of evidence and rationality varying across clusters—rendering a claim credible in one context but illegitimate in another. Without a reflexive analysis of how justification functions across these boundaries, CP risks conflating coexistence with equivalence, treating incommensurable epistemologies as if they were symmetrical. This leads to a form of implicit relativism that confuses epistemology—how knowledge is justified—with ontology—what exists independently of that knowledge [48,49]. Absent clear criteria for distinguishing warranted knowledge from misinformation or strategic claims, the very standards of epistemic justification risk being eroded.

Indeed, the embodied and intuitive heuristics underpinning commonsense knowledge can expose the limits of scientism and prompt critical reflection on how epistemic authority is constructed and mobilized to serve political or private interests. Yet, reliance solely on intuitively salient cues can also distort judgment, with serious consequences for science-informed decision-making - particularly in sensitive domains such as public health [27,50]. In such contexts, appeals to “multiple truths” may function less as genuine expressions of epistemic pluralism than as strategic instruments to obscure evidence, manufacture doubt, or delay policy action. Rather than fostering dialogical engagement between epistemologies, these dynamics often manifest as struggles over credibility and legitimacy, where tensions among knowledge, trust, and power shape both counter-authoritative narratives and the conditions under which scientific authority itself is produced and sustained.

In this context, conspiratorial epistemology can be understood as emerging from a dialectical tension between epistemic mistrust and biased information processing [51]. The former is not simply a cognitive distortion but is often rooted in epistemic injustice - the systematic exclusion of certain groups from recognition as credible knowers [52]. When grounded in experiences of exclusion or institutional failure, such mistrust may be epistemically justifiable, functioning as a form of situated common sense that responds to structural inequities in credibility. It can motivate the search for alternative explanatory frameworks, a process with the potential to challenge dominant epistemic hierarchies.

However, in fragmented and low-credibility information environments—particularly under post-truth conditions—this search becomes increasingly vulnerable to distortion and misinformation. When trustworthy epistemic resources are inaccessible, individuals may default to unreliable accounts as compensatory meaning-making strategies. Limited source-evaluation ability [53], together with motivated reasoning and identity-protective cognition, further amplifies susceptibility to misleading narratives [54]. Over time, these dynamics can reconstruct epistemic mistrust, transforming it from skepticism toward specific content into broader institutional mistrust - casting doubt on science governance as a legitimate source of epistemic authority.

This reconstructed epistemic mistrust is frequently filtered through conspiracy narratives, when instrumentalized, transform skepticism into indiscriminate institutional mistrust. While experiences of injustice can render such mistrust understandable, they do not automatically legitimize the claims built upon it. Conspiracy theories are “for losers” [55], as they often emerge among individuals who feel politically marginalized or powerless, channeling frustration toward perceived out-groups [56] - particularly when access to reliable information or cognitive resources is limited. These narratives are frequently amplified by strategic actors who exploit mistrust to undermine epistemic authority, often without substantive justification. [57]. Science-related populism exemplifies this dynamic, valorizing everyday common sense while portraying scientific expertise as detached, elitist, or untrustworthy [58]. This epistemic antagonism is consistently associated with declining confidence in science, even when controlling for education, demographics, and religiosity [59].

Investigating this contested epistemic space confronts a dilemma: the Scylla of equating epistemic dominance with objectivity - thus naturalizing existing credibility hierarchies - and the Charybdis of treating claims from marginalized groups as inherently legitimate. Both positions obscure the relational and contextual processes through which epistemic claims gain credibility. A more robust standard for epistemic justification lies not in presumed objectivity or authenticity, but in whether knowledge practices broaden the social and institutional conditions for inclusive, reflexive, and progressive inquiry.

Accordingly, it is important to distinguish between critical epistemic mistrust - grounded in reflective engagement with scientific authority - and the dogmatic embrace of counter-narratives. Collapsing the public’s diverse modes of engaging with science into simple pro-anti categories obscures the complexity of how people position themselves toward scientific institutions. Orientations toward science develop through the interaction of cultural values, reasoning styles, and institutional trust [20], which together shape responses that are multidimensional and domain-specific [60]. As Godin and Gingras [61] note, *scientific culture* comprises both individual and social dimensions - it involves personal knowledge and attitudes but also the cultural and institutional contexts that confer meaning upon science. Conspiratorial reasoning, therefore, can express either critical questioning or closed-minded rejection, depending on its epistemic grounding and social context.

### C. Reassessing the Knowledge–Conspiracy–Trust Pathway: Beyond the Deficit Model and Cognitive Polyphasia

Although the multidimensional structure of public understanding of science has been widely examined [22,62,63], less attention has been given to the mediating pathways linking its components. Recent studies have begun to address this gap by identifying indirect relations through which education, values, and cognitive factors interact with scientific trust. Research indicates that education shapes conspiratorial thinking and trust in science through cognitive and value-based mediators [64,65], and that generalized faith in science mediates the effects of religious orthodoxy and conspiracy beliefs on mistrust [66]. Similarly, right-wing authoritarianism predicts pseudoscientific and paranormal beliefs indirectly through social axioms [67], while beliefs about scientific authority and the roles of scientists and citizens are mediated by epistemic trust and deference to expertise [68]. In the health domain, vaccine conspiracy beliefs mediate the relationship between scientific knowledge and vaccine hesitancy, operating through reduced trust in medical science [69,70].

A key limitation in existing research on mediating pathways is the tendency to conflate science-related conspiracism with either institutional mistrust or epistemic deficit, treating them as interchangeable indicators of disengagement from science. This study differentiates these orientations by defining science knowledge as a measure of epistemic competence, conspiracism as an epistemic orientation - a distinct way of constructing and legitimizing knowledge- and institutional mistrust as an attitudinal stance toward scientific authority. Although these dimensions frequently correlate [67, 71], they interact in complex ways that warrant closer conceptual separation.

Conspiratorial reasoning operates as an epistemic framework for interpreting uncertainty and perceived injustice through assumptions of hidden intent and systemic deception, while institutional mistrust reflects the evaluative stance arising from this filtered view of scientific authority. Conspiracists often frame their pursuit as a rational, truth-seeking endeavor, positioning themselves as investigators uncovering hidden realities through critical inquiry [72]. As a mediating heuristic, it fills gaps in factual understanding, expressing both a desire for coherence and the constraints of epistemic vulnerability in complex information environments [73]. In this capacity, conspiratorial reasoning can act either as a form of epistemic agency or as a source of distortion, reinterpreting evidence according to alternative criteria of legitimacy when epistemic trust is strained. It is thus neither the inverse of scientific literacy nor a mere expression of institutional mistrust, but a distinct epistemic mode emerging from epistemic mistrust under conditions of uncertainty, positioned along a continuum between reflective engagement and dogmatic closure [74].

Adding to this complexity, the boundary between factual knowledge and evaluative preconceptions is often blurred, a challenge reflected in how the construct is operationalized. Standardized measures of scientific knowledge in surveys frequently lack theoretical grounding and psychometric clarity, showing low discriminatory power, modest reliability, and pronounced ceiling effects, while remaining vulnerable to acquiescence bias and statistical artifacts [75]. These instruments tend to conflate epistemic, attitudinal, and cultural dimensions, interpreting variations in trust or interpretation as knowledge deficits. Many items are culturally inflected, capturing moral and institutional judgments as much as cognitive understanding—in effect, measuring perceptions of authority and institutional mistrust rather than knowledge alone [76]. The result is an implicit fuzziness masked by explicit linearity, as complex epistemic distinctions collapse into a single continuum of correctness.

A similar pattern characterizes measures of science attitudes. Classical models distinguish between perceptions of science’s benefits and reserves about its societal consequences [62,77]. More recent approaches build on this distinction by further differentiating views of science as a driver of technological progress from its role as a moral or institutional authority [59,78]. Rather than existing on a single continuum, these dimensions are analytically orthogonal, delineating a two-dimensional landscape in which people can support scientific progress while expressing reservations about its ethical or institutional impact.

Beyond their multidimensionality and fuzziness, science-related constructs are also embedded in macro influences. Broader contexts shape how science and its institutions are interpreted, extending beyond factual knowledge alone [79]. According to research, the knowledge-attitude relationship varies across national settings - stronger in intermediate economies and weaker in less developed or post-industrial contexts - reflecting how different configurations activate distinct predictors of attitudes [80–82]. However, multilevel evidence is mixed: while regional and national socioeconomic factors account for substantial variation beyond individual knowledge and attitudes in the EU [83], broader cross-national analyses find more limited cultural differences [84]. Regarding conspiracy beliefs, they are shaped by both individual (e.g., scientific knowledge) and contextual factors (e.g., national affluence or regime type) shape, with education and vaccine attitudes exerting stronger effects in wealthier democracies [85], where even those with limited knowledge or higher mistrust of scientists are less prone to endorse science-related conspiracy theories than their counterparts in more authoritarian settings [86]. Overall, higher average knowledge or educational investment appears associated with *context-dependent, complex dynamics* that interweave multiple constructs rather than yielding uniform improvements in trust. [87]

In conclusion, the relationship between knowledge, conspiratorial reasoning, and science mistrust is best understood as a context-dependent configuration in which individual epistemic and motivational factors are embedded within broader sociocultural conditions. These dynamics determine how knowledge translates into conspiratorial reasoning and, through it, into institutional mistrust across levels of analysis.

### D. Hypotheses and Model Overview

Building on this framework, this study adopts a middle-range theoretical approach [13,88] that integrates these conceptual insights into an empirical multilevel structural model. This enables us to move beyond Deficit and Contextualist paradigms by examining how science knowledge and related covariates influence institutional mistrust indirectly via conspiratorial reasoning, and how this pathway is contingent on informational, motivational, and sociocultural conditions.

Scientific and conspiratorial epistemologies represent distinct but interconnected expressions of the same underlying epistemic domain. Contestations of scientific authority are further differentiated into two interrelated components: conspiratorial reasoning, an *epistemic* orientation, and institutional mistrust, an *attitudinal* stance. The former spans a continuum from flexible, case-specific questioning to rigid, generalized suspicion, reflecting differences in how individuals construct and evaluate knowledge claims. The latter, by contrast, captured through the *reserve construct*, reflects a broader normative attitude against science encompassing concerns about institutional trust, epistemic authority, and the societal role of science.

Positioned at the intersection of epistemic and attitudinal domains, conspiracy reasoning operates as a bridge between how individuals *know* and how they *trust*. This intermediary position justifies its specification as a mediating mechanism linking epistemic orientations to attitudinal outcomes. Modeling conspiratorial reasoning as the antecedent of mistrust is consistent with longitudinal evidence showing that counter-authoritative beliefs more reliably predict subsequent declines in trust in experts, institutions, and science-related behaviors than the reverse [89]. The model further incorporates three clusters of covariates - Cognitive and Informational Factors, Perceptual and Science-Related Beliefs, and Ideological and Sociocultural Factors - to account for additional sources of variation influencing both conspiracy beliefs and mistrust.

> **H1.** The indirect effect of science knowledge on mistrust through conspiratorial reasoning will be significant and stronger than corresponding indirect effects for the covariates.

Beyond the core mediational pathway, and consistent with our contingency framework, the model incorporates moderated mediation and mediated moderation, in which knowledge × covariate interactions shape and transmit effects on mistrust through conspiratorial reasoning.

> **H2a.** The knowledge → conspiracy → mistrust pathway will be conditioned by informational, perceptual, ideological, and sociocultural variables.

While individual epistemic capacities establish a baseline for the relationship between knowledge and attitudes, their effects are shaped by broader cultural and political contexts [12,61]. Religiosity, in particular, represents a salient contextual factor influencing how scientific knowledge relates to mistrust through conspiratorial reasoning. The literature remains divided: some studies emphasize incompatibilities between religious and scientific epistemologies—especially concerning claims to epistemic authority [19,90,91] - whereas others find that religious commitment and scientific understanding can coexist within individuals and communities [41].

Building on this mixed evidence, we propose that the conspiracy-mediated link between scientific knowledge and mistrust is contingent on the broader religious climate that defines epistemic legitimacy and evaluative norms, reflecting neither a universal deficit nor a coexistence pattern.

> **H2b.** The mediated pathway between individual knowledge and mistrust will be conditioned by macro-level religiosity, such that its strength and direction depend on the prevailing religious climate.

At the macro level, the model extends the individual-level mediation framework to test whether similar relationships emerge among aggregated constructs, without committing ecological fallacy. Specifically, it examines whether regions with higher average science knowledge show lower prevalence of conspiratorial orientations and, in turn, reduced mistrust. This captures how collective scientific understanding and shared epistemic climates reflect broader cognitive-informational, political-cultural, and socioeconomic conditions shaping societal orientations toward science and authority.

> **H3**. The indirect effect of science knowledge on mistrust through conspiratorial reasoning will be significant and stronger than corresponding indirect effects for the covariates at the macro level as well.

## III. Data and Methods

### A. Data

Data were drawn from Eurobarometer 95.2 (ZA7782), a cross-national survey conducted in April–May 2021 across 38 European countries. The target population comprised residents aged 15 and older, selected via multistage, stratified random sampling by region and population size. Interviews were administered using computer-assisted personal (57%) and web interviewing (43%) modes. The initial dataset included 37,079 individuals in 308 regions. After excluding cases with >20% missing attitude or >30% missing knowledge responses, the final analytic sample comprised 35,075 participants.

Missing data were handled via multiple imputation using chained equations (MICE) with 12 auxiliary covariates representing demographics, science engagement, political orientation, and paradata.

### B. Measures

#### Latent Variables

All latent constructs were estimated using multidimensional item response theory (MIRT), which provides item-level parameters—difficulty, discrimination, and, for dichotomous items, guessing—along with factor scores that incorporate measurement error (see Supplementary materials), suitable for multilevel structural equation modeling (MSEM). Reliabilities for latent variables were in the good range. The full wording of all items used in the analysis(Appendix A) and MIRT results (Appendix B) are available in Supplementary materials.

#### Science Knowledge (Knowledge) and Conspiracy Reasoning (Conspiracy)

An 11-item battery assessed scientific knowledge and conspiratorial reasoning using true–false items adapted from established science literacy instruments. To avoid conflating ignorance with uncertainty, “Don’t know” responses were treated as missing and imputed following Mondak [92]. Exploratory MIRT revealed three dimensions: Basic Literacy (e.g., basic school knowledge), Scientific Knowledge (e.g., understanding of physical and biological mechanisms), and Conspiratorial Reasoning (e.g., beliefs about hidden scientific agendas, suppressed cures, or doubts about scientific consensus).

Two items on human evolution and climate change cross-loaded on both knowledge and conspiracy factors, consistent with the framework treating factual understanding and epistemic contestation as distinct yet intersecting orientations.The conspiracy factor represents an epistemic continuum from reflective questioning of uncertainty to dogmatic closure, where stronger alignment across items indicates generalized suspicion toward scientific authority. This dual loading illustrates the continuum’s logic: low knowledge responses denote factual ignorance, whereas high alignment within the conspiratorial factor reflects an epistemic style of closure rather than a simple knowledge deficit. Final scores were derived from the confirmatory analysis, in which the Basic Literacy factor was excluded due to low reliability (ω = .35).

#### Science Attitudes-Reserve (Mistrust)

A 19-item battery assessed attitudes toward science on a 5-point Likert scale. Exploratory MIRT revealed three dimensions: Reserve (7 items reflecting mistrust of science and scientists), Promise (9 items reflecting optimism about science’s problem-solving potential), and Deference (3 items reflecting respect for scientific authority). Reserve and Promise replicated prior findings, confirming their cross-population robustness. Only Reserve was retained as the key indicator of mistrust, capturing skepticism against scientific authority, including concerns about scientists’ power, accountability, trustworthiness, and the societal implications of scientific progress.

#### Science Engagement (Engagement)

A 12-item battery assessed science-related participation on a 4-point frequency scale. Exploratory MIRT identified two dimensions: Active Engagement (8 items; e.g., signing petitions, participating in clinical trials) and Passive Engagement (4 items; e.g., discussing science with friends, watching documentaries). Only Passive Engagement was retained due to floor effects in Active Engagement and its stronger theoretical relevance to knowledge and conspiracy.

#### Negative Scientist Perception

This construct comprised 10 binary items (e.g., *narrow-minded, arrogant, immoral*). Positive trait items were reverse-coded so that higher scores indicated greater attribution of negative characteristics.

#### Unequal Benefits of Science and Technology (Unequal Benefits)

This construct included three items (e.g., *“Science mostly improves the lives of the well-off”*), refined from an eight-item battery, rated on a five-point Likert scale.

#### Observed Variables

Single-item predictors captured populist decision-making, religiosity, right political orientation, use of social media as the main source of science information (social media), worldview (cooperative vs. threat-oriented), information about scientific discoveries (information), and perceived inefficacy of science (Perceived vaccine inefficacy; reverse-coded evaluations of vaccines and disease prevention).

#### Control Variables

Control variables included socio-demographics—age, respondent and parental education, and economic strain (difficulty paying bills)—in line with standard practices in the conspiracy literature [93] (Smallpage et al., 2020), as well as paradata [85], such as interview mode (CAPI vs. CAWI), ‘don’t know’ propensity (share of DK responses), and guessing propensity (ratio of incorrect to incorrect plus DK responses)

#### Level 2 (Regional) Variables

Regional variables included both aggregated individual constructs and external contextual indicators. Regional means were computed automatically within the lavaan MSEM framework for Conspiracy, Mistrust, Knowledge, Unequal Benefits, Negative Scientist Perceptions, Religiosity, and Satisfaction with Democracy.

Regional electoral integrity perception index was modeled as a latent construct using an eight-item, four-point EVS/WVS 2017–2022 battery assessing electoral fairness and institutional integrity (Supplementary materials), while trust toward people of another nationality (single observed item) was drawn from the same source; both were aggregated by region.

Contextual indicators included per capita purchasing power standard (PPS) and the gender employment gap [94], indexing regional economic development and gender inequality. Because the EVS/WVS dataset did not cover Belgium, Ireland, Luxembourg, and Malta, regional scores for Electoral Integrity and Outgroup Trust in these countries were imputed based on regional covariates (PPS, university enrollment, disposable income, gender employment gap, and high-tech employment).

### C. Use of AI Tools

Artificial intelligence tools were used to assist with language editing, stylistic refinement, and support in R coding. These tools served solely as auxiliary aids; all substantive analyses, coding logic, interpretation of results, and theoretical development were independently conducted and verified by the authors.

## IV. Results

### A. Analysis

Multilevel structural equation models were estimated in lavaan package [95], decomposing variance into within- and between-EU region components. Random intercepts were modeled via latent between-level variables.

Intraclass correlations confirmed substantial regional clustering (ICC = .31 for *Conspiracy*; ICC = .21 for *Mistrust*). A Hausman-style correlated random effects test indicated endogeneity of random effects (Wald χ²(1) = 893.71, *p* < .001) and significant within–between differences (Wald χ²(2) = 303.25, *p* < .001), supporting the MSEM approach to decompose variance across levels.

Models were estimated using maximum likelihood with the NLMINB optimizer. Because *lavaan* does not yet support random slopes, all slopes were fixed across regions.

Model fit was assessed using standard indices [(Kline, 2023)], indicating good overall fit: CFI = .95, TLI = .87, RMSEA = .025, SRMR*_within_* = .01, and SRMR*_between_*= .075.

Indirect effects were tested using Monte Carlo confidence intervals to account for non-normality. Explained variance for all endogenous variables: *Conspiracy* (*R²_within_* = .67; *R²_between_* = .84) and *Mistrust* (*R²_within_*= .58; *R²_between_* = .83).

Interaction terms were computed externally from centered variables. Level-1 predictors were group-mean centered to capture within-region effects, and corresponding regional means were grand-mean centered to represent between-region effects. All Level-2 contextual variables were z-standardized.

All covariates described in the *Measures* section were included at their respective levels to adjust for demographic, methodological, and contextual influences.

### B. Descriptive Statistics and Correlations

Tables 1 and 2 present means, standard deviations, and zero-order correlations for individual-level (Level 1) and regional-level (Level 2) variables. At the individual level, knowledge is moderately negatively correlated with conspiracy (r = −.48) and mistrust (r = − .39), while conspiracy is positively associated with mistrust (r = .48). A similar pattern appeared at the regional level: average knowledge showed strong negative correlations with both conspiracy (r = −.89) and mistrust (r = −.81), and conspiracy were strongly correlated with mistrust (r = .85). Correlations among other variables at both levels ranged from weak to moderate

**Table 1.**
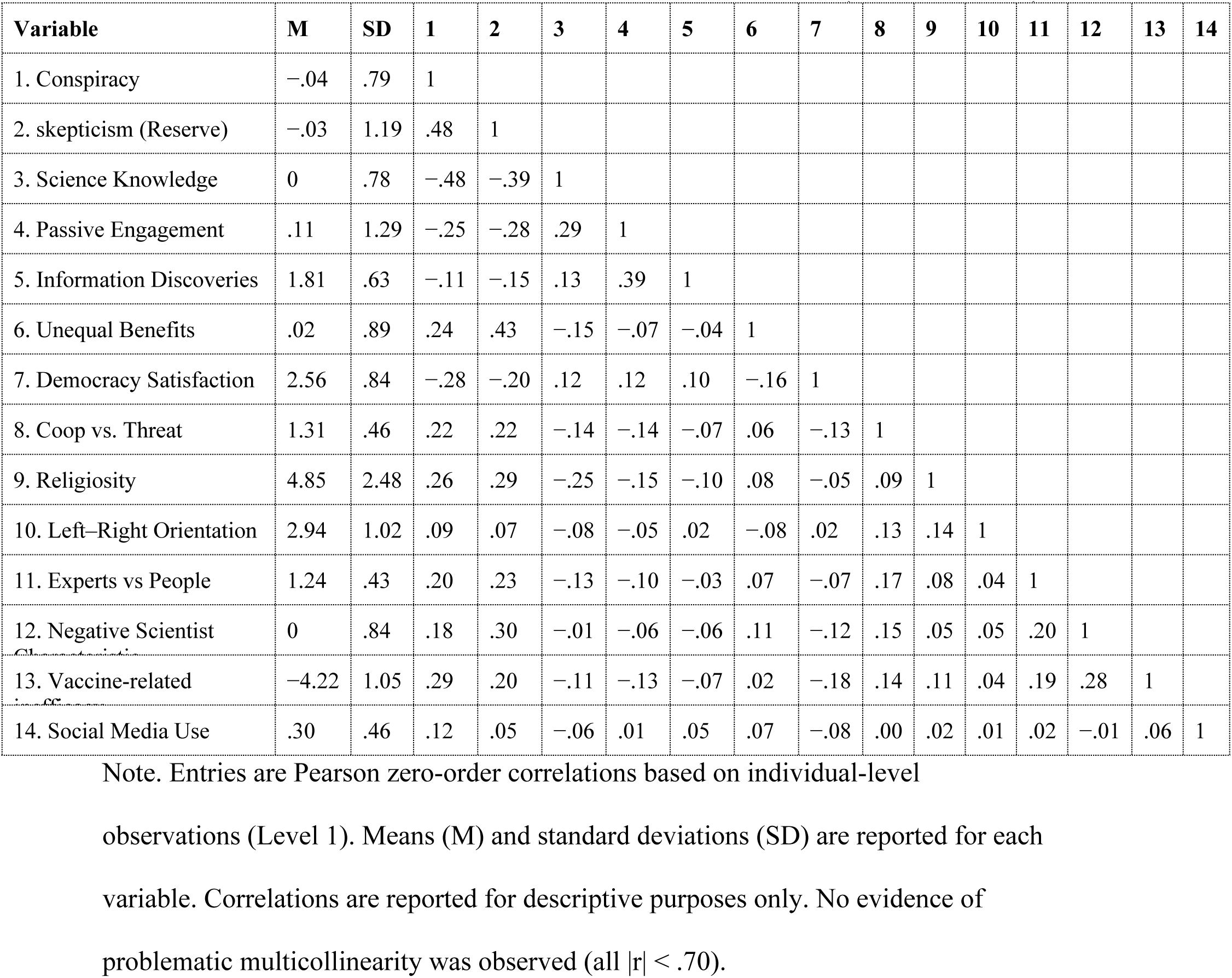
Means, Standard Deviations, and Zero-Order Correlations (Individual Level)

**Table 2.** Means, Standard Deviations, and Zero-Order Correlations among Regional Level Variables.

| Variable | M | SD | 1 | 2 | 3 | 4 | 5 | 6 | 7 | 8 | 9 | 10 | 11 | 12 |
| --- | --- | --- | --- | --- | --- | --- | --- | --- | --- | --- | --- | --- | --- | --- |
| 1. Conspiracy beliefs | -0.04 | 0.44 | — |  |  |  |  |  |  |  |  |  |  |  |
| 2. skepticism (reserve) | -0.03 | 0.57 | .85 | — |  |  |  |  |  |  |  |  |  |  |
| 3. Science knowledge | 0.00 | 0.41 | -.89 | -.81 | — |  |  |  |  |  |  |  |  |  |
| 4. Democracy satisfaction | 2.56 | 0.36 | -.64 | -.54 | .51 | — |  |  |  |  |  |  |  |  |
| 5. Unequal benefits of science | 0.02 | 0.30 | .56 | .77 | -.52 | -.48 | — |  |  |  |  |  |  |  |
| 6. Religiosity | 4.85 | 1.14 | .73 | .66 | -.70 | -.39 | .42 | — |  |  |  |  |  |  |
| 7. Cooperation vs. threat worldview | 1.31 | 0.13 | .57 | .56 | -.48 | -.40 | .31 | .44 | — |  |  |  |  |  |
| 8. Democratic trust (z) | 0.00 | 1.00 | -.78 | -.60 | .68 | .57 | -.33 | -.67 | -.47 | — |  |  |  |  |
| 9. Trust in other nations (z) | 0.00 | 1.00 | .74 | .62 | -.63 | -.59 | .39 | .61 | .46 | -.62 | — |  |  |  |
| 10. Negative scientist characteristics | 0.00 | 0.27 | .02 | .09 | .12 | .05 | -.09 | .03 | .24 | -.02 | .11 | — |  |  |
| 11. Expert vs. people orientation (z) | 0.00 | 1.00 | -.69 | -.56 | .62 | .55 | -.35 | -.48 | -.44 | .58 | -.52 | .07 | — |  |
| 12. Gender equality gap (z) | 0.00 | 1.00 | .58 | .50 | -.57 | -.40 | .35 | .61 | .40 | -.55 | .53 | -.08 | -.41 | — |

**Notes.** Entries are Pearson correlations based on regional aggregates (Level 2; N =
35,075). Means (M) and standard deviations (SD) are reported for each variable. All
correlations are statistically significant at $p < .001$ due to large sample size unless547 otherwise noted. No evidence of problematic multicollinearity was observed (all $|r| <$

### C. Within Level (Individual) Effects

#### Main Mediation Path: Knowledge → Conspiracy → Mistrust

Scientific knowledge was strongly and negatively associated with conspiracy beliefs (*β* = −.57, *p* < .001). Its direct effect on mistrust was weaker (*β* = −.16, *p* < .001) than the corresponding bivariate correlation (*r* = −.39), consistent with prior research showing attenuation once confounding variables are controlled (see Table 3).

**Table 3.**
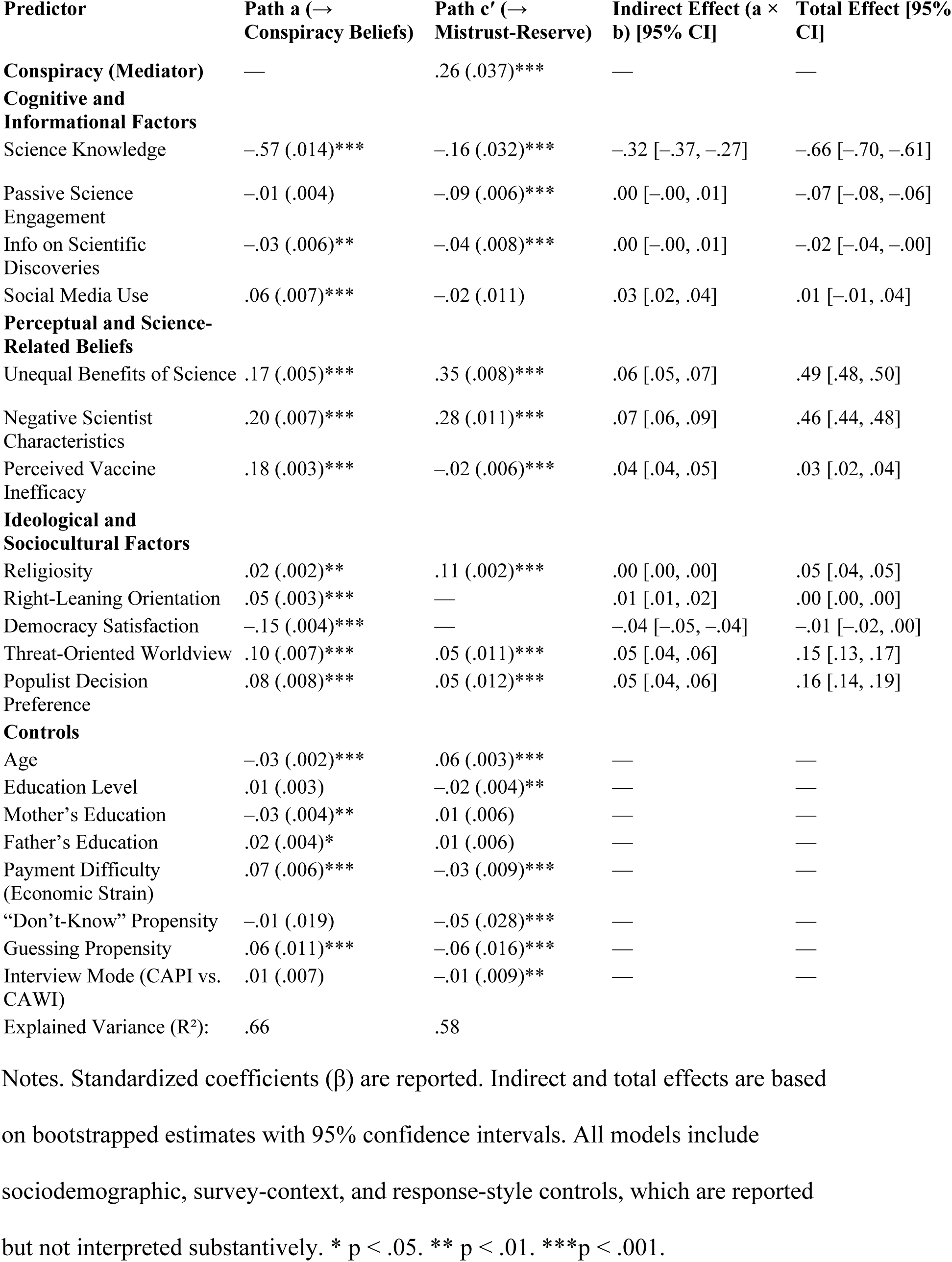
Standardized Path Coefficients (β), Standard Errors (SE), and Indirect/Total Effects for the Within-Level Model.

Knowledge exerted a significant indirect effect on mistrust via conspiracy beliefs (*β* = −.32, 95% CI [−.37, −.27]), accounting for nearly half of the total effect (*β* = −.66, 95% CI [−.70, −.61]). These findings support conspiracy reasoning as the principal mediating mechanism linking scientific knowledge to institutional mistrust.

#### Cognitive and Informational Variables

Engagement with scientific content was unrelated to conspiracy reasoning and influenced mistrust only through a direct pathway. Exposure to scientific information reduced conspiracy beliefs but exhibited offsetting direct and indirect effects on mistrust, resulting in a negligible total effect. Social media use was positively related to conspiracy beliefs and affected mistrust primarily through this indirect pathway, yielding a modest yet significant mediated effect.

#### Perceptual and Science-Related Predictors

Perceived unequal benefits of science and negative views of scientists were both positively associated with conspiracy beliefs and mistrust, showing weak indirect but stronger direct effects. These predictors produced some of the largest total effects in the model, although mediation was weaker. Perceived vaccine inefficacy positively predicted conspiracy beliefs but had a weak negative direct effect on mistrust, producing a small net positive total effect after accounting for indirect pathways.

#### Ideological and Sociocultural Factors

Right-leaning ideology and satisfaction with democracy predicted mistrust indirectly through conspiracy beliefs, with smaller direct effects. Religiosity, perceived societal threat, and populist orientations demonstrated both direct and mediated associations, indicating partial mediation. Although their indirect effects were weaker than those for knowledge, they nonetheless contributed to total effects on mistrust.

#### Moderated Mediation

Moderated mediation models (Hayes, Model 15) indicated that both engagement and self-perceived information about scientific discoveries significantly strengthened the indirect effect of knowledge on mistrust through conspiracy beliefs (indices of moderated mediation = −.027 and −.028, respectively; see Table 4). Conditional estimates showed that the indirect and total effects of knowledge were more negative at higher levels of these moderators. Religiosity and perceived unequal benefits of science also moderated the mediation: a positive index for religiosity (.010) suggested attenuation of the negative indirect effect at higher religiosity, whereas a negative index for unequal benefits (−.014) indicated that perceived inequality amplified the indirect effect.

**Table 4.** Knowledge-Conditioned Interaction Effects (mediated moderation and moderated mediation-Hayes 15)

| Moderator | Interaction Coefficients ($\beta$) | Indirect Pathway | Conditional Indirect Effect (Low / High) [95% CI] | Conditional Total Effect (Low / High) [95% CI] |
| --- | --- | --- | --- | --- |
| **Moderated mediation (Hayes Model 15)** |  |  |  |  |
| Passive Science Engagement | K×Eng: $-.025^{***}$; C×Eng: $.045^{***}$ | **Index** = $-.027$ [$-.035$, $-.019$] | Low: $-.29$ [$-.34$, $-.24$] / High: $-.35$ [$-.41$, $-.30$] | Low: $-.60$ [$-.64$, $-.55$] / High: $-.72$ [$-.76$, $-.67$] |
| Self-Perceived Information | K×Info: $-.047^{***}$; C×Info: $.046^{***}$ | **Index** = $-.028$ [$-.042$, $-.013$] | Low: $-.30$ [$-.35$, $-.24$] / High: $-.35$ [$-.40$, $-.29$] | Low: $-.59$ [$-.63$, $-.54$] / High: $-.72$ [$-.77$, $-.68$] |
| Religiosity | K×Rel: $.010^{**}$; C×Rel: $-.017^{***}$ | **Index** = $.010$ [$.006$, $.014$] | Low: $-.34$ [$-.40$, $-.29$] / High: $-.30$ [$-.35$, $-.25$] | Low: $-.70$ [$-.74$, $-.65$] / High: $-.61$ [$-.65$, $-.57$] |
| Unequal Benefits of Science | K×UB: $-.034^{***}$; C×UB: $.024^{**}$ | **Index** = $-.014$ [$-.024$, $-.004$] | Low: $-.31$ [$-.36$, $-.25$] / High: $-.33$ [$-.39$, $-.28$] | Low: $-.62$ [$-.66$, $-.57$] / High: $-.70$ [$-.74$, $-.65$] |
| **Mediated moderation** |  |  |  |  |
| Negative Scientist Characteristics | Conspiracy: $-.016^{**}$; Mistrust: $.053^{***}$ | **ab** = $-.009$ [$-.015$, $-.002$] | **Indirect** = $-.009$ [$-.015$, $-.002$] | **Total** = $.045$ [$.028$, $.062$] |
| Regional Religiosity (Level 2) | Conspiracy: $.018^{**}$; Mistrust: $.067^{***}$ | **Index** = $.010$ [$.005$, $.015$] | Low: $-.33$ [$-.39$, $-.28$] / High: $-.31$ [$-.36$, $-.26$] | Low: $-.74$ [$-.79$, $-.69$] / High: $-.57$ [$-.61$, $-.53$] |

**Notes.** The first row reports mediated moderation; remaining rows report moderated mediation (Hayes Model 15). Conditional effects are evaluated at $\pm 1$ SD (or low vs.
high regional religiosity). All effects use bootstrapped 95% confidence intervals. $p < .05$ , $p < .01$ , $p < .001$ .

#### Mediated Moderation

Mediated moderation analyses indicated that the interaction between scientific knowledge and negative perceptions of scientists was negatively associated with conspiracy beliefs and positively associated with mistrust, producing a small but significant indirect effect via conspiracy beliefs (ab = −.009, 95% CI [−.015, −.002]) and a positive total effect (β = .045, 95% CI [.028, .062]).

#### Cross-Level Moderation

At the regional level, the indirect effect of individual knowledge on mistrust via conspiracy beliefs varied by regional religiosity (index = .010). Conditional analyses revealed that the negative indirect effect of knowledge was stronger in less religious regions and diminished as regional religiosity increased, resulting in a weaker total effect under more religious contexts.

### D. Between-Level (Regional) Effects

At the regional level, knowledge was strongly and negatively associated with both conspiracy beliefs and mistrust, producing a sizable indirect effect (−.130, 95% CI [−.190, −.099]) and a substantial direct effect on mistrust (*β* = −.334, *p* < .001).

Together, these yielded a large total effect (−.570, 95% CI [−.673, −.470]), indicating that regions with higher knowledge levels exhibited lower overall mistrust (see Table 5, Panel A).

**Table 5.** Between-Level (Regional) Mediation and Cross-Level Moderated Mediation Effects.

| Predictor | Path a → Conspiracy Beliefs (β, SE) | Path c' → Mistrust (β, SE) | Indirect Effect (a×b) [95% CI] | Total Effect [95% CI] |
| --- | --- | --- | --- | --- |
| **Conspiracy Beliefs (Mediator)** | — | .209 (.075)** | — | — |
| **Cognitive & Perceptual Contexts** |  |  |  |  |
| Science Knowledge | -.472 (.038)*** | -.334 (.062)*** | -.130 [-.209, -.056] | -.570 [-.673, -.470] |
| Unequal Benefits of Science | .111 (.033)*** | .516 (.046)*** | .036 [.013, .067] | .841 [.752, .931] |
| Negative Scientist Perceptions | .108 (.035)*** | .205 (.054)*** | .037 [.012, .068] | .368 [.262, .474] |
| **Political & Ideological Contexts** |  |  |  |  |
| Democracy Satisfaction | -.150 (.035)*** | .007 (.047) | -.046 [-.082, -.018] | -.036 [-.126, .055] |
| Electoral Integrity | -.116 (.016)*** | .011 (.021) | -.014 [-.028, -.005] | -.008 [-.050, .035] |
| Religiosity | .091 (.013)** | .137 (.017)*** | .009 [.002, .019] | .074 [.040, .107] |
| Threat-Oriented Worldview | .065 (.081)* | .101 (.106)*** | .050 [.007, .109] | .424 [.214, .636] |
| Out-Group Distrust | .129 (.014)*** | -.042 (.019) | .015 [.005, .028] | -.008 [-.045, .029] |
| **Economic Contexts** |  |  |  |  |
| Purchasing Power | -.081 (.015)* | -.052 (.020) | -.010 [-.022, -.002] | -.042 [-.081, -.002] |
| Gender Employment Inequality | .022 (.012) | -.119 (.015)*** | .002 [-.004, .009] | -.053 [-.083, -.024] |

Panel B. Cross-Level Moderated Mediation by Regional Negative Scientist Perceptions
| Moderator (Level 2) | Interaction Effect on Reserve (β) | Indirect Pathway (Index of Moderated Mediation) [95% CI] | Conditional Indirect Effect (Low / High) [95% CI] | Conditional Total Effect (Low / High) [95% CI] |
| --- | --- | --- | --- | --- |
| Negative Scientist Perceptions (Regional) | **K×SciNeg:** -.151***;<br>**C×SciNeg:** -.200*** | .332 [.199, .560] | Low: -.220 [-.250, -.124]<br>/ High: -.040 [-.073, .053] | Low: -.530 [-.417, -.352]<br>/ High: -.610 [-.448, -.513] |

**Notes.** Panel A reports between-level (regional) mediation effects via conspiracy beliefs. Panel B reports cross-level moderated mediation of the science knowledge → skepticism association by regional negative perceptions of scientists, including the index of moderated mediation and conditional indirect and total effects evaluated at ±1 SD of the moderator. All confidence intervals are bootstrapped 95%. $p < .05$ , $p <$ $.01$ , $p < .001$ .

Unequal benefits of science and negative perceptions of scientists showed similar but weaker patterns. Both were positively related to conspiracy beliefs and mistrust, yielding significant indirect effects (.036, 95% CI [.013, .067]; .037, 95% CI [.013, .069]) and strong direct effects, resulting in substantial total effects.

Political and cultural factors displayed mixed associations. Democracy satisfaction and electoral integrity each reduced conspiracy beliefs and produced significant negative indirect effects on mistrust (−.046, 95% CI [−.082, −.018]; −.014, 95% CI [−.028, −.005]) but negligible direct effects. In contrast, religiosity positively predicted conspiracy beliefs and mistrust, yielding a small indirect effect (.009, 95% CI [.002, .019]) and a substantial direct effect (*β* = .137, *p* < .001). Threat-oriented worldviews and out-group distrust similarly increased conspiracy beliefs, generating modest indirect effects (.015, 95% CI [.005, .028]) but weak or negative direct effects on mistrust, producing minimal total influence.

Economic variables showed modest associations. Higher purchasing power predicted fewer conspiracy beliefs, yielding a small negative indirect effect on mistrust (−.010, 95% CI [−.022, −.002]) without a significant direct effect. Gender employment gap had no indirect effect but was directly and negatively associated with mistrust (*β* = −.119, *p* < .001), producing a small total effect driven by the direct pathway.

#### Between-Level Moderated Mediation by Regional Negative Scientist Perceptions

Between-level moderated mediation analyses revealed that the indirect link between regional knowledge and mistrust via conspiracy beliefs varied with regional perceptions of scientists (see Table 5, Panel B). Both the knowledge × negative perceptions (*β* = −.151, *p* < .001) and conspiracy × negative perceptions (*β* = −.200, *p* < .001) interactions were significant, with a corresponding index of moderated mediation of .332 (95% CI [.199, .560]). Conditional estimates showed that the negative indirect effect of knowledge on mistrust was stronger in regions with more positive views of scientists and attenuated where negative perceptions were higher.

## V. Discussion

### A. Within (individual) level structural pathways Central Mediation Path (H1)

Consistent with H1, knowledge stands out as the predictor for which the indirect effect via conspiracy beliefs is both significant and substantially larger than that of any covariate, accounting for a sizable share of its total association with mistrust. Specific covariate effects along this pathway are as follows:

Informational antecedents exhibit divergent pathways to mistrust. Engagement and self-perceived information reduce mistrust primarily through direct mechanisms. In contrast, social media use fosters conspiracy thinking, which indirectly heightens mistrust—though its total effect remains modest. This finding is consistent with findings that conspiracy-oriented individuals disengage from conventional information channels [96] in favor of informal networks such as social media [30]. These environments can amplify conspiratorial narratives and facilitate the diffusion of misinformation, particularly among individuals predisposed to epistemic insecurity, motivated reasoning and within cultural climates characterized by uncertainty avoidance [97, 98].

Perceptual variables reveal that mistrust differs according to its object. Perceived unequal benefits of science and negative views of scientists exert primarily direct effects resonating with Goldenberg’s [99] argument that much “science denial” reflects institutional rather than epistemic mistrust. In such cases, mistrust arises as an attitudinal response to perceived institutional injustice or as heightened vigilance toward authority [100], expressing a normative critique through which knowledge influences mistrust directly, without mediation by conspiratorial reasoning. By contrast, the mediation observed for vaccine efficacy suggests that rejecting biomedical consensus often relies on the explanatory coherence of conspiracy theories—to resolve the tension between heuristic insecurity and established evidence [101]. Together, these patterns reveal two distinct modes of resistance to science: one rooted in *critical mistrust*, which engages in normative and institutional critique, and another characterized by *conspiratorial closure*, which resolves epistemic uncertainty by constructing rigid alternative frameworks that offer coherence—albeit through distorted or unfalsifiable explanations.

Among ideological and sociocultural factors, religiosity shapes science mistrust primarily through direct normative judgments rather than via conspiratorial mediation. Although often correlated, religiosity and conspiracy thinking typically serve as competing frameworks for interpreting power and epistemic authority, especially in polarized contexts [102]. Mistrust emerges when science is seen as encroaching on domains—such as morality—where religion claims primacy [59]. In contrast, right-wing orientation and satisfaction with democracy influence mistrust mainly through conspiratorial pathways, despite their modest total effects—consistent with findings that link these factors to conspiracy belief [103–106].

This pattern aligns with research showing that right-wing ideology is more consistently associated with conspiracy thinking, whereas religiosity more directly predicts science mistrust [19]. Threat-based worldviews and populist preferences exert the strongest combined influence, highlighting how perceived threat and anti-expert sentiment promote conspiratorial reasoning and, through it, science mistrust [107]. Although rarely examined as mediators, the centrality of conspiracy in our model echoes evidence that such epistemologies link political discontent, institutional distrust, and opposition to scientific authority [108].

#### Conditional Pathways and Contextual Moderation

After discussing the structural pathways, we next consider how the knowledge-conspiracy-mistrust path is conditioned by informational, normative, and perceptual contexts, reflecting both moderated mediation and mediated moderation dynamics.

#### Moderated Mediation (H2a)

Informational moderators—engagement and perceived information—condition the indirect effect of knowledge within the mediation pathway. When high, they enhance knowledge’s capacity to constrain conspiratorial interpretations; when low, this effect weakens. Yet under limited knowledge, elevated engagement or perceived information may instead amplify conspiracy’s influence on mistrust by fostering overconfidence without accuracy [78, 109]. Acting as epistemic amplifiers, these moderators intensify the conspiratorial pathway rather than offsetting weak epistemic competence.

For cultural and ideological moderators, religiosity and perceived inequality shape the mediation pathway in distinct yet complementary ways. Religiosity operates in the abstract moral domain, weakening the protective effect of knowledge by filtering trust through faith-based norms rather than empirical reasoning. Mistrust in highly religious contexts thus reflects adherence to alternative moral frameworks rather than conspiratorial cognition. In contrast, conspiratorial reasoning becomes more salient among less religious individuals with limited knowledge, functioning as a secular-epistemic substitute for faith-based meaning-making. This aligns with work portraying conspiracism as a *secular religion* that offers cognitive closure and certainty to those lacking epistemic resources [110]. Religiosity therefore acts as both a normative filter and a competing system of moral interpretation.

Perceived inequality, by contrast, is rooted in concrete experiential intuition. It weakens knowledge’s ability to constrain conspiratorial reasoning while intensifying its translation into mistrust. Here, knowledge remains epistemically valid but loses normative force when fairness is questioned. Among high-knowledge individuals, this tension manifests as critical disengagement grounded in perceived institutional injustice; among low-knowledge individuals, limited epistemic resources and lived experiences of unfairness jointly foster conspiratorial interpretations [50]. Compared to religiosity’s abstract moral grounding, perceptions of inequality derive from tangible social experience, serving not as an alternative moral framework but as an affective trigger that reactivates conspiratorial meaning-making when institutional fairness is doubted.

#### Mediated Moderation (H2b)

In the mediated moderation between knowledge and negative perceptions of scientists, conspiracy beliefs function as a diminished translation mechanism, as skepticism shifts away from alternative explanatory narratives toward actor-focused credibility evaluations. When scientific actors are perceived as untrustworthy, skepticism can be justified without recourse to conspiratorial explanations (Vaupotič et al., 2021 [111]). Importantly, negative perceptions of scientists do not imply rejection of scientific knowledge but rather reconfigure the basis of skepticism, making it plausible that skepticism is grounded in distrust of scientific representatives rather than in epistemic opposition to science itself. This pattern is consistent with the *science confidence gap* observed particularly among less-educated groups, for whom skepticism reflects institutional distrust rather than epistemic rejection (Achterberg et al., 2017 [112]).

#### Cross-Level Moderation (H2c)

At the cross-level, regional religiosity acts as a normative climate shaping the individual link between knowledge and mistrust. In highly religious regions, the protective effect of scientific knowledge weakens, reflecting a partial decoupling of epistemic competence from trust in science. Here, mistrust is driven less by conspiratorial reasoning than by faith-based evaluative frameworks. Conversely, in less religious regions—especially among low-knowledge individuals—mistrust becomes more contingent, relying on conspiratorial narratives to resolve epistemic uncertainty in the absence of strong normative anchors. These patterns suggest that the relationship between science and religion reflects neither a simple coexistence nor a deficit logic but an *interactional dynamic*, in which the epistemic and normative domains condition one another depending on the broader religious climate.

Taken together, the effects of ideological and cultural factors reveal two distinct logics shaping the mediation pathway. In religious contexts, knowledge is reshaped by metaphysical commitments that place faith above epistemic authority. By contrast, under institutional conditions like perceived inequality, mistrust emerges through critical engagement, where knowledge is challenged as part of a broader sense of injustice.

### B. Between level (regional) structural path (H3)

At the between level, analyses corroborate H3.Across predictors, regional effects consistently operate by modifying the strength or relevance of the knowledge–conspiracy–mistrust pathway rather than replacing it, underscoring its role as the primary organizing structure of regional science mistrust. At the regional level, the outcome reflects institutional mistrust rather than individual skepticism, emphasizing shared evaluative climates rather than personal doubt. Specific covariate indirect and total effects along this pathway are discussed below.

Regions characterized by greater inequality in the perceived benefits of science and more negative views of scientists exhibit higher overall mistrust—both directly and indirectly through conspiracy climates. The coexistence of strong direct and mediated effects indicates that mistrust climates embody shared normative evaluations of scientific institutions, operating within and beyond conspiratorial belief systems.

Conspiracy climates translate structural inequality into epistemic mistrust, linking perceived unfairness to the collective erosion of institutional legitimacy. These dynamics mirror broader processes through which unequal distributions of knowledge, authority, and resources corrode trust at both epistemic and normative levels [113].

Parallel to individual and cross-level findings, regional religiosity shows a predominantly direct association with mistrust, with only a small secondary indirect pathway via regional conspiracy beliefs, indicating that regional value climates ground mistrust primarily through broader faith-based orientations rather than conspiratorial sense-making.

Political legitimacy shapes institutional mistrust primarily by constraining conspiracy belief, aligning with prior research [103,114]. Regions with higher democratic satisfaction and stronger electoral integrity show lower levels of conspiracism, though their direct impact on mistrust remains limited. However, this relationship should be interpreted with caution, as democracy is strongly collinear with other structural variables such as economic development, institutional effectiveness, and corruption control [86]. The negligible total effects suggest that democracy acts more as a contextual buffer—structuring the epistemic environment in which scientific claims are evaluated—than as a direct determinant of science-related attitudes [115].

Threat-oriented worldviews and out-group distrust relate to science mistrust through distinct mediating pathways. In contexts of heightened threat sensitivity, mistrust reflects a broader climate of insecurity, where conspiracy beliefs amplify but do not fully account for mistrust toward science. In contrast, out-group distrust contributes to science mistrust primarily when organized into conspiratorial narratives; intergroup suspicion alone does not translate into mistrust without such framing. These patterns highlight conspiracy thinking as a selective mediator: it channels out-group distrust into science mistrust by combining it with broader epistemic suspicion, while threat-based mistrust emerges more independently. This aligns with research showing that conspiracy theories, particularly under climates of insecurity, foster general institutional distrust by portraying decision-making processes as fundamentally unfair and illegitimate—even when the conspiracy claims are not directly tied to the institutions in question [116,117].

Regional economic conditions show relatively weaker and uneven associations. Higher purchasing power is associated with lower regional conspiracy beliefs and modestly reduced mistrust, suggesting that economic security constrains conspiratorial sense-making [118,85] without directly anchoring science attitudes. By contrast, gender employment inequality is directly linked to lower regional mistrust but shows no meaningful mediation via conspiracy beliefs, indicating that some structural inequalities shape mistrust through non-conspiratorial pathways. Overall, economic contexts appear to function as background conditions [117] rather than primary drivers of regional science mistrust.

The moderated mediation analysis indicates that the indirect association between regional scientific knowledge and institutional mistrust via regional conspiracy beliefs is stronger in regions where scientists are viewed more positively. In these contexts, lower average knowledge is associated with stronger conspiracy climates and, in turn, higher mistrust, whereas higher knowledge constrains conspiratorial interpretations.

By contrast, in regions where scientists are viewed negatively, this pathway is attenuated and mistrust is anchored more directly in normative evaluations of scientific actors rather than in regional knowledge distributions. This pattern suggests that macro-level legitimacy—scientific or political—functions as an epistemic buffer: in high-legitimacy settings, knowledge and trust remain more closely aligned, while in contexts marked by generalized elite distrust, scientists become targets of broader institutional skepticism [115]. Under such conditions, conspiratorial narratives may coexist with institutional cynicism, but mistrust is increasingly grounded in direct evaluations of scientific actors rather than mediated through conspiracy beliefs [120].

## VI. Conclusion & Limitations

Taken together, our findings position scientific knowledge neither as a uniformly protective factor, as assumed by deficit models, nor as evidence of stable coexistence between incommensurable epistemologies, as implied by cognitive polyphasia. Instead, knowledge functions as a contingent epistemic resource whose effects depend on informational, normative, and institutional contexts. Its capacity to attenuate mistrust is more direct in settings that support engagement, perceived competence, and the epistemic authority of science, but becomes increasingly mediated by conspiracy thinking where motivated cognition, perceived injustice, or epistemic mistrust shape evaluations of science.

In this vein, science mistrust reflects neither simple ignorance nor entrenched epistemic plurality, but contextually modulated pathways through which knowledge, conspiracy thinking, and mistrust are configured. By tracing these pathways across individual and subnational levels, our analysis moves beyond broad paradigm debates to specify how science mistrust is structured by individual characteristics and across diverse political and cultural settings. In doing so, it bridges the explanatory clarity of deficit-oriented accounts with the contextual sensitivity of constructivist approaches, advancing a middle-range synthesis in which the epistemic authority of science is negotiated through the interplay of cognitive resources, sociocultural meanings, and institutional credibility.

These insights, however, should be viewed as provisional given several empirical and methodological constraints. First, the analyses rely on cross-sectional and secondary data, which limits causal inference and the ability to align measurement precisely with the theoretical constructs under investigation. Although the models are theoretically grounded and temporally plausible, longitudinal and experimental designs are needed to capture dynamic feedback processes between knowledge acquisition, conspiratorial beliefs, and mistrust. Second, while regional-level aggregation enhances statistical power and allows more granular multilevel analysis than national comparisons, regional associations describe shared contextual climates rather than individual attitudes and should not be interpreted as direct analogues of individual-level processes, raising the usual cautions regarding ecological inference. At the same time, regional aggregation may obscure localized meaning systems and interpretive practices through which science skepticism is constructed. This underscores the value of mixed-method and qualitative approaches capable of tracing how knowledge, trust, and conspiratorial beliefs are negotiated in specific social contexts. Third, although the models incorporate a broad set of cognitive, normative, and sociopolitical moderators, other influences such as media ecologies, elite signaling, or issue-specific scientific controversies may further condition these pathways.

Within this framework, our contribution lies not in offering definitive claims but in providing a provisional snapshot of dynamic pathways, one that invites future research into feedback mechanisms, situated epistemic-cultural practices, and alternative mediating influences.

## Acknowledgments

The authors gratefully acknowledge the support of the Bulgarian National Science Fund for funding the project “Images of Science in the Digital World: Between Social Networks and Internet Media” (Contract No. КП 06-Н55/6, November 2021), and the Institute of Philosophy and Sociology at the Bulgarian Academy of Sciences as the base organization. This research was also supported by TÜBİTAK under project number 220N219, “Science News in Bulgarian and Turkish Media: Automated Monitoring of Science Culture Indicators Across Time and Cultures,” and by the Caring Communities project (DRP0200240), co-funded by the European Union through the Interreg Danube Region Programme.

## Supporting information

S1 File. Supplementary tables and model parameters.

## Data Availability

Data used in this study are publicly available from the GESIS Data Archive (Eurobarometer 95.2, ZA7782).

